# Inference of super-exponential human population growth via efficient computation of the site frequency spectrum for generalized models

**DOI:** 10.1101/022574

**Authors:** Feng Gao, Alon Keinan

## Abstract

The site frequency spectrum (SFS) and other genetic summary statistics are at the heart of many population genetics studies. Previous studies have shown that human populations had undergone a recent epoch of fast growth in effective population size. These studies assumed that growth is exponential, and the ensuing models leave unexplained excess amount of extremely rare variants. This suggests that human populations might have experienced a recent growth with speed faster than exponential. Recent studies have introduced a generalized growth model where the growth speed can be faster or slower than exponential. However, only simulation approaches were available for obtaining summary statistics under such models. In this study, we provide expressions to accurately and efficiently evaluate the SFS and other summary statistics under generalized models, which we further implement in a publicly available software. Investigating the power to infer deviation of growth from being exponential, we observed that decent sample sizes facilitate accurate inference, e.g. a sample of 3000 individuals with the amount of data expected from exome sequencing allows observing and accurately estimating growth with speed deviating by 10% or more from that of exponential. Applying our inference framework to data from the NHLBI Exome Sequencing Project, we found that a model with a generalized growth epoch fits the observed SFS significantly better than the equivalent model with exponential growth (*p*-value = 3.85 × 10^−6^). The estimated growth speed significantly deviates from exponential (*p*-value << 10^−12^), with the best-fit estimate being of growth speed 12% faster than exponential.

## Introduction

Summary statistics of genetic variation play a vital role in population genetics studies, especially inference of demographic history. In particular, the site frequency spectrum (SFS) is a vital summary statistic of genetic data and is widely utilized by many demographic inference methods applied to humans and other organisms (Marth et al. 2004; Gutenkunst et al. 2009; Excoffier et al. 2013; Bhaskar et al. 2015; Liu and Fu 2015). Other demographic inference methods are based on sequentially Markov coalescent and utilize the most recent common ancestor (*T*_MRCA_) and linkage disequilibrium patterns (Li and Durbin 2011; Harris and Nielsen 2013; MacLeod et al. 2013; Sheehan et al. 2013; Schiffels and Durbin 2014). As another example, several studies used the average pairwise difference between chromosomes (Hammer et al. 2008; Gottipati et al. 2011; Arbiza et al. 2014) and the SFS (Keinan et al. 2009) to study the relative effective population sizes between the human X chromosome and the autosomes. The wide application of such genetic summary statistics stresses the need for their fast and accurate computation under any model of demographic history, instead of their estimations via simulations or approximations (e.g., Hudson 2002; Gutenkunst et al. 2009).

Several recent demographic inference studies showed evidence that human populations have undergone a recent epoch of fast growth in effective population size (Gutenkunst et al. 2009; Coventry et al. 2010; Gravel et al. 2011; Nelson et al. 2012; Tennessen et al. 2012; Gazave et al. 2014). However, the above studies assumed that the growth is exponential. The observation of huge amount of extremely rare, previously unknown variants in several sequencing studies with large sample sizes (Nelson et al. 2012; Tennessen et al. 2012; Fu et al. 2013) and the recent explosive growth in census population size suggests that human population might have experienced a recent super-exponential growth, i.e. growth with speed faster than exponential (Coventry et al. 2010; Keinan and Clark 2012; Reppell et al. 2012, 2014). Hence, recent studies presented a new generalized growth model that extends the previous exponential growth model by allowing the growth speed to be exponential or faster/slower than exponential (Reppell et al. 2012, 2014). Modeling the recent growth by this richer family of models holds the promise of a better fit to human genetic data, and can also be applicable to other organisms that experienced growth. However, only simulation approaches are currently available for evaluating such a generalized growth demographic model (Reppell et al. 2012), which makes inference of demographic history computational intractable.

In this study, we first provide a set of explicit expressions for the computation of five summary statistics under a model of any number of epochs of generalized growth: (1) the time to the most recent common ancestor (*T*_MRCA_), (2) the total number of segregating sites (*S*), (3) the site frequency spectrum (SFS), (4) the average pairwise difference between chromosomes per site (*π*), and (5) the burden of private mutations, BPM (*α*), a summary statistic that has been recently introduced as sensitive to recent growth (Keinan and Clark 2012; Gao and Keinan 2014). We introduce a new software package that implements these expressions, EGGS (Efficient computation of Generalized models’ Genetic summary Statistics), which facilitates fast and accurate generation of these summary statistics. We show that the numerically computed summary statistics match well with simulation results, and facilitates computation that is orders of magnitudes faster than that of simulations. By performing demographic inference on the SFS generated from simulated sequences, we then explore how many samples are needed for recovering parameters of a recent generalized growth epoch. Finally, we applied the software to investigate the nature of the recent growth in humans by inferring demographic models using the SFS of synonymous variants of 4,300 European individuals from the NHLBI Exome Sequencing Project (Tennessen et al. 2012; Fu et al. 2013).

## Methods and Materials

### Generalized demographic models

A demographic model *N*(*T*) describes the changes of effective population size *N* against time *T*. We consider time, measured in generations, as starting from 0 at present and increasing backward in time. Furthermore, we consider the families of demographic models that are constituted by any number of epochs of generalized growth, along the lines of (Bhaskar and Song 2014). More formally, there exists a minimal positive integer *L* such that the demographic history of a population can be split into a model with *L* + 1 epochs that are split by *L* ordered different time points *T*_1_, *T*_2_, …,*T*_*L*_ (*T*_0_ = 0 < *T*_1_ < *T*_2_ *…* < *T*_*L*_ < *T*_*L*+1_ = ∞), with the *k*^th^ epoch starting from *T*_*k–1*_ and lasting through *T*_*k*_ (thus the last epoch starts at time *T*_*L*_ and continues into indefinite past, *T*_*L+1*_ = ∞). Such a history is considered as a generalized model if the population size in each epoch *N*(*T*_*k*–1_ ≤ *T* < *T*_*K*_) can be described by the following differential equation regarding time *T* (Reppell et al. 2012, 2014):

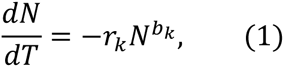

where *k* = 1, 2, …, *L* + 1. Each epoch (*k* = 1, *2*, *…*, *L* + 1) can hence capture variety of changes patterns in effective population size. Specifically, if *r*_*k*_ = 0, this epoch is of constant population size. When *r*_*k*_ ≠ 0, *b*_*k*_ controls the growth speed of this epoch: (1) if *b*_*k*_ = 1, the epoch is of exponential growth (*r*_*k*_ > 0) or decline (*r*_*k*_ < 0) with rate *r*_*k*_; (2) if *b*_*k*_ > 1, the epoch is of faster-than-exponential (super-exponential) growth (*r*_*k*_ > 0) or decline (*r*_*k*_ < 0); (3) if *b*_*k*_ < 1, the epoch is of slower-than-exponential (sub-exponential) growth (*r*_*k*_ > 0) or decline (*r*_*k*_ < 0). Linear growth or decline is also a special case of generalized models when *b*_*k*_ = 0. An illustration of a generalized model with 5 epochs is provided in Figure 1, and more detailed explanation and illustrations in the Supporting Information.

**Figure 1.**
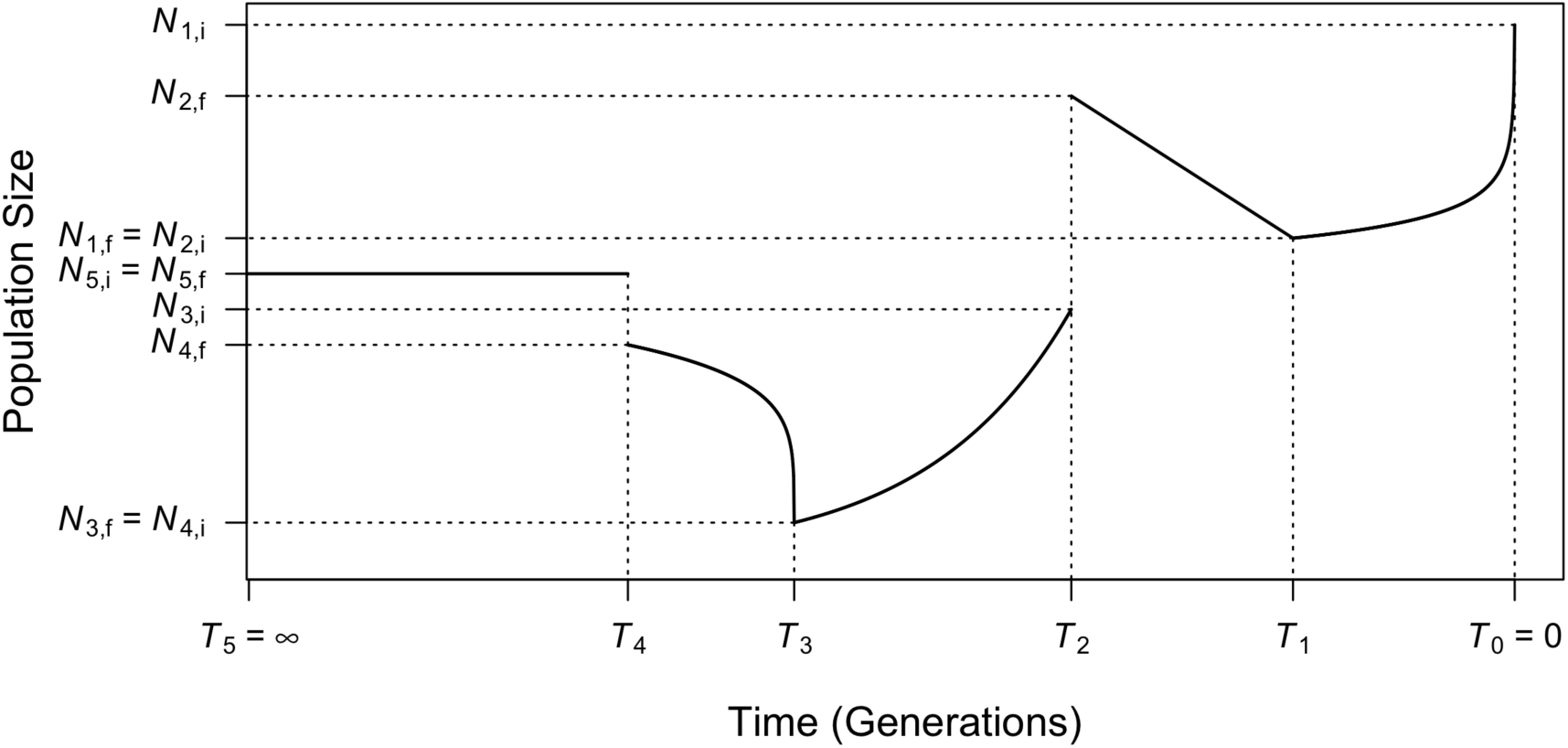
Illustration of an example of a generalized demographic model as introduced in the first section of Methods and Materials. This model consists of 5 epochs (starting from present): (1) Faster-than-exponential (*b* > 1) growth (looking forward in time) from *N*_1,f_ to *N*_1,i_ between *T*_0_ = 0 and *T*_1_; (2) Linear decline (a special case of generalized growth, *b* = 0) from *N*_2,f_ to *N*_2,i_ between *T*_1_ and *T*_2_; (3) Exponential growth (a special case of generalized growth, *b* = 1) from *N*_3,f_ to *N*_3,i_ between *T*_2_ and *T*_3_; (4) Slower-than-exponential (*b* < 1) decline from *N*_4,f_ to *N*_4,i_ between *T*_3_ and *T*_4_; (5) Constant population size (a special case of generalized growth, *r* = 0) at *N*_5,i_ = *N*_5,f_ starting from *T*_4_ and lasts indefinitely backward in time (*T*_5_ = ∞). The ending population size of the previous epoch is not necessarily the beginning population size of the next epoch (e.g., *N*_2,f_ ≠ *N*_3,i_, *N*_4,f_ ≠ *N*_5,i_), corresponding to an instantaneous population size change at that time.

The solution to Equation (1) has been derived (Reppell et al. 2012, 2014) to be

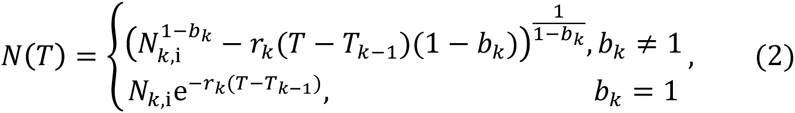

where *N*_*k*,*i*_ is the initial population size of the *k*^th^ epoch. Each epoch *k* is defined by 4 parameters: the starting population size (*N*_*k*,*i*_), the ending population size (*N*_*k*,*f*_), the duration of the epoch (*T*_*k*_ − *T*_*k*–1_) and the growth speed parameter *b*_*k*_. The growth rate parameter, *r*_*k*_ is an immediate function of these parameters, *r*_*k*_ = *r*_*k*_ (*N*_*k*,*i*_, *N*_*k*,*i*_, *b*_*k*_, *T*_*k*_ – *T*_*k*–1_), and hence does not need to be provided as an independent variable in defining the changes in effective population size during an epoch. Note that *N*_*k*+1,i_, the starting population size of the (*k* + 1)^th^ epoch is not necessarily the same as *N*_*k*,*f*_, the ending population size of the *k*^th^ epoch. When it is not, it entails an instantaneous change in population size at time *T*_*k*_.

### Explicit expressions for summary statistics of demographic models under arbitrary population size functions

Previous studies showed that under Kingman’s coalescent, given a demographic model *N*(*T*), the expected time of the most recent common ancestor 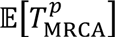 can be calculated by (Polanski et al. 2003)

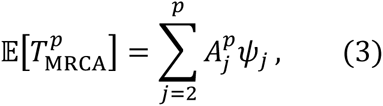

where the superscript *p* is the number of chromosomes (twice the diploid sample size), *Ψ*_*j*_ is the expected time of the first coalescent event when there are *j* chromosomes at present, which is given by

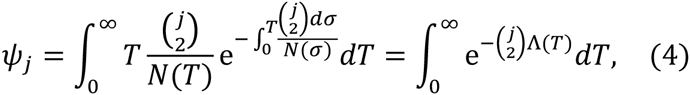

where 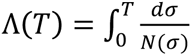, and 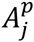 are constants (Tavare 1984; Takahata and Nei 1985; Polanski et al. 2003) provided in Supporting Information.

The expected full relative SFS 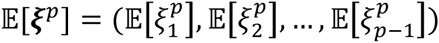 can be computed by the following set of equations (Polanski and Kimmel 2003):

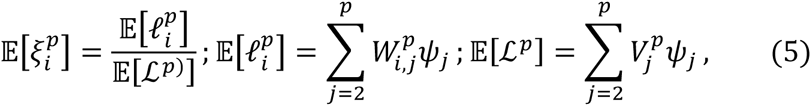

where 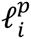is the length of branches in the genealogy that have *i* descendants and 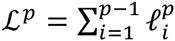 is the total length of all branches in the coalescent tree. The quantities 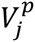 and *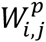* are constants (Polanski and Kimmel 2003), which we provide in the Supporting Information.

Naturally, the expected number of segregating sites is given by

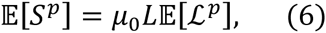

where *μ*_0_ is the mutation rate per site per generation and *L* is the length of the locus under consideration. The average pairwise difference between chromosomes per site, 𝔼[ π], can be calculated by

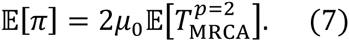

The expected burden of private mutations (*α*) at sample size of 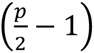, defined as the proportion of heterozygous sites in a new diploid individual that are homozygous in the previous 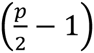 individuals, 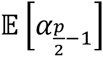 can be approximated by (Gao and Keinan 2014)

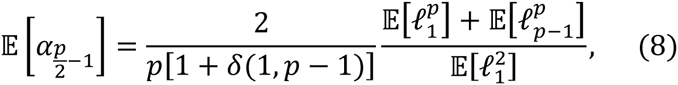

where *δ*(*x*, *y*) is Kronecker delta function.

The detailed description of the five summary statistics mentioned above is included in the Supporting Information.

### Evaluation of the expected time to the first coalescent event under generalized models

The core of the evaluation lies in finding feasible and numerically stable functions for calculating *Ψ*_*j*_, the expected time of the first coalescent event when there are *j* chromosomes at present. Previous studies give explicit expressions of *Ψ*_*j*_ for a demographic model constructed by exponential and constant-size epochs (Polanski and Kimmel 2003; Bhaskar et al. 2015). In this study, we give a comprehensive set of formulas for generalized models as introduced above. Define 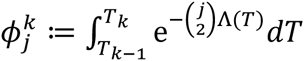 then 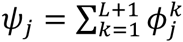, where (*L* + 1) is the total number of epochs. The quantity 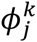 can be computed numerically by the following set of equations:

(1) If *r*_k_ = 0:

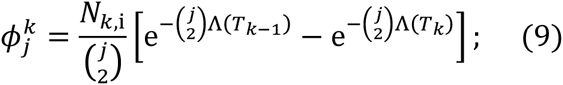

(2) If *b*_k_ = 0, *r*_k_ ≠ 0:

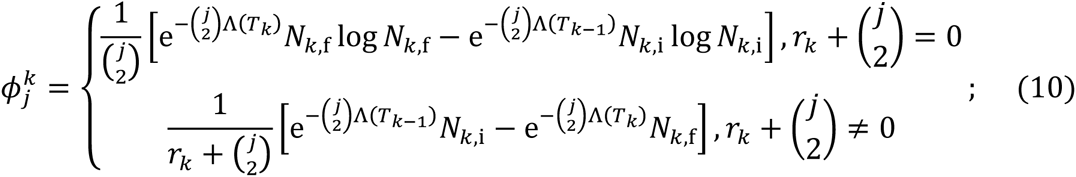

(3) If *b*_k_ > 0 and *r*_k_ > 0, or *b*_k_ = 1 and *r*_k_ < 0;

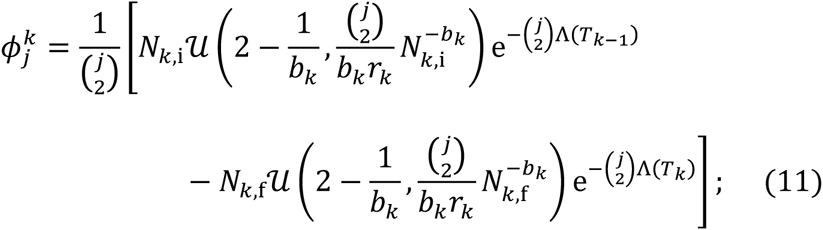

(4) If *b*_k_ < 0 and *r*_k_ > 0:

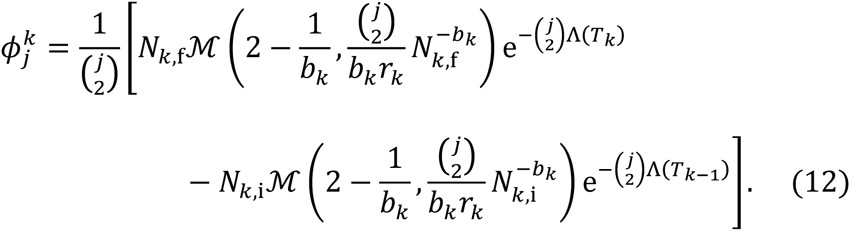

The expressions of the Λ(*T*) function are given in the Supporting Information. The function 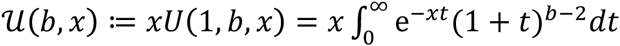, where *U*(*a*, *b*, *x*) is the confluent hypergeometric function of the second kind (Gradshteĭn et al. 2007). The function 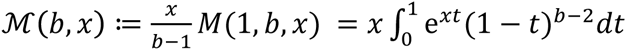, where *M*(*a*, *b*, *x*) is the confluent hypergeometric function of the first kind (Gradshteĭn et al. 2007). The exponential growth or decline then becomes a special case of *u*(*b*, *x*) when *b* = 1 and *x* ≠ 0,

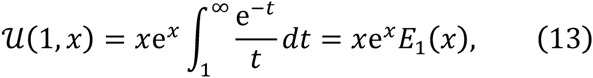

where *E*_1_ (*x*) is the Exponential Integral (Gradshteĭn et al. 2007), which has been shown by previous studies (Polanski and Kimmel 2003; Bhaskar et al. 2015). We could not find feasible and numerically stable closed-form formulas for 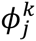 when the population size decreases forward in time in a manner that is not linear or exponential (i.e. *r*_*k*_ < 0 and *b*_*k*_ ∉ {0,1}). In these scenarios, we used Gauss-Legendre quadrature (Kahaner et al. 1988) for efficient numerical evaluation of relevant functions (see Supporting Information for a detailed description).

### Software implementation

The above expressions are implemented in a software package, EGGS (Efficient computation of Generalized models’ Genetic summary Statistics). The software and source code are publicly available from our website (http://keinanlab.cb.bscb.cornell.edu). Compiled versions for Linux and MAC OS are available. Source code is in C++, with no external libraries needed for compilation. Additional information of implementation is included in the Supporting Information and in the manual that accompanies the software online.

### Demographic models assumed in this study

The demographic models used in this study are based on the inferred European history presented by Gazave *et al*. 2014 (Figure 2, in black), which contains two bottlenecks (Keinan et al. 2007) and a recent exponential growth epoch. Specifically, Gazave *et al.* 2014 model has a constant effective population size of 10,000 (diploid) individuals before 4,720 generations ago, and goes through the ancient bottleneck between 4,720 generations ago and 4,620 generations ago with a population size of 189. The population size then recovers to 10,000 diploids until 720 generations ago, from which time the recent bottleneck starts with a size of 549. At 620 generations ago, the population size recovers to 5,633 individuals. The recent growth epoch starts from 140.8 generations ago and lead to population size of 654,000 at present. The parameters of the original recent growth epoch were varied to incorporate generalized growth effects.

**Figure 2.**
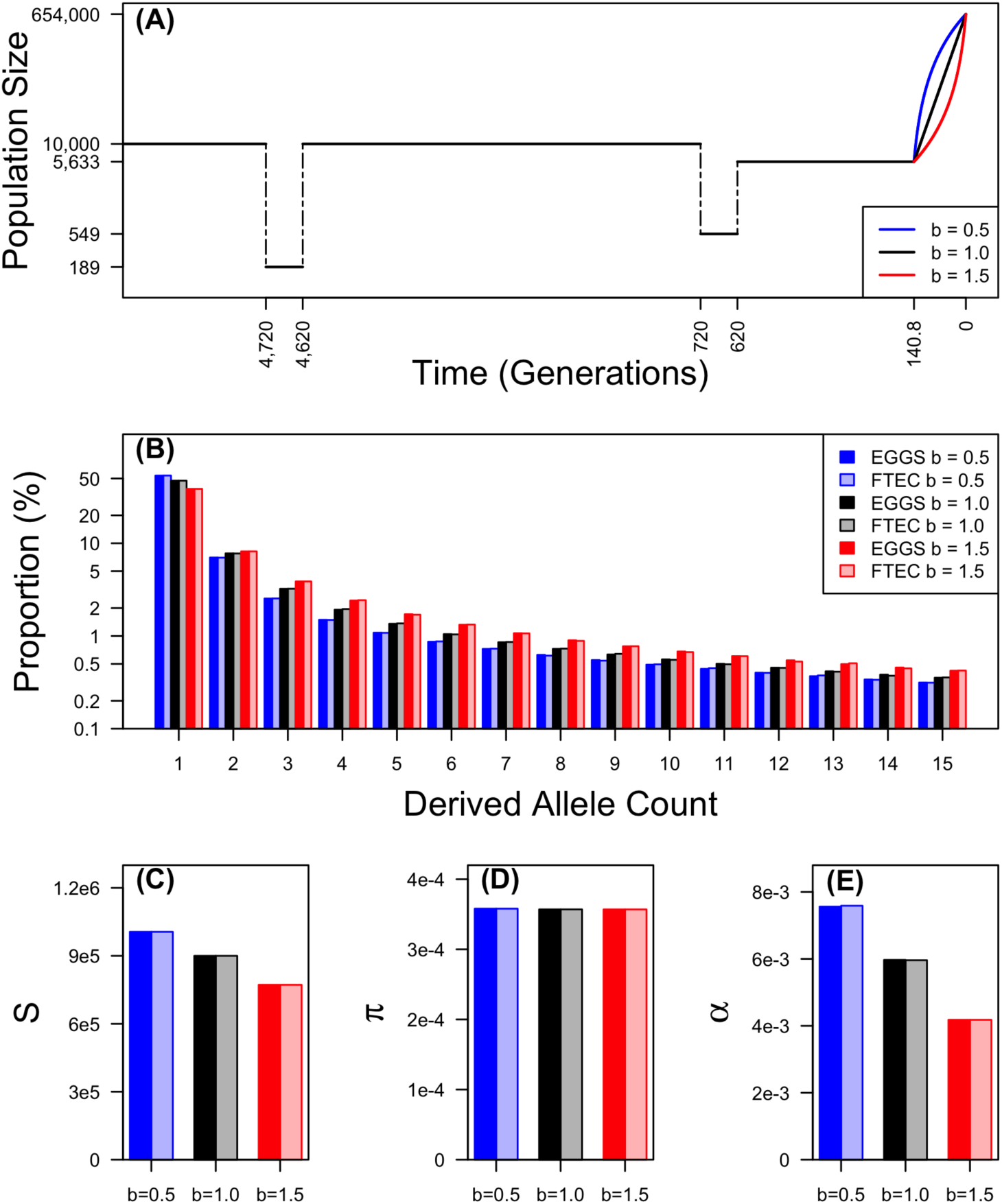
(A) Demonstration of the demographic models considered for evaluating the accuracy of our calculations as implemented in EGGS (first section of Results). This two-bottleneck model has the same population size and time throughout history as in the inferred European history in Gazave *et al*. 2014, with the exception that we varied the growth speed parameter of the recent growth epoch to be *b* = 0.5 (sub-exponential, blue), *b* = 1.0 (exponential as in Gazave *et al*. 2014, black) and *b* = 1.5 (super-exponential, red). The *y*-axis shows effective population size of diploid individuals on a log-scale. (B)-(E) The comparison of the first 15 entries of the SFS (B), the total number of segregating sites (*S*) across all 200,000 loci (1,000 bp-long each) (C), the expected pairwise difference between chromosomes per base pair (D) and the burden of private mutation (*α*) as the percentage of heterozygous variants in one individual that are monomorphic in the rest of the sample of 999 individuals (E) computed numerically in EGGS (dark bars) and simulated by FTEC (light bars) for the demographic models shown in (A): *b* = 0.5, blue; *b* = 1.0, black; *b* = 1.5, red, with a sample size of 1,000 individuals (2,000 chromosomes). The *y-axis* in (B) is on log scale.

### Optimization method of demographic inference based on the site frequency Spectrum

Demographic inference in this study was based on the observed allele frequency counts from the simulated or real dataset. To determine the fitness of a model to the observed data, we calculated the composite log likelihood by

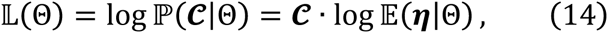

where ***ℂ*** is a vector of the observed folded allele frequency counts and 𝔼(*η*|Θ) is the computed folded SFS under demographic model Θ. More detailed description can be found in the Supporting Information.

To search for the maximum likelihood point over the parameter space, we applied the ECM method (Meng and Rubin 1993; Excoffier et al. 2013). 100 ECM cycles were performed for each run of inference. We obtained 95% confidence intervals of parameter estimates via block bootstrapping of the data 200 times.

### Processing of NHLBI Exome Sequencing Project data for demographic history inference

The NHLBI Exome Sequencing Project (ESP) data (Tennessen et al. 2012; Fu et al. 2013) contains deep sequencing of 4,300 individuals of European ancestry. An important feature of these data is the high sequencing coverage, which allows to capture accurately very rare variants, which constitutes the part of the SFS that is most enriched for information on recent population growth (Keinan and Clark 2012; Tennessen et al. 2012; Gao and Keinan 2014). To reduce the effect of selection as much as possible while keeping sufficient amount of data, we chose to use the SFS calculated from synonymous single nucleotide variants (SNVs) only, as previously performed by Tennessen *et al*. 2012. To further improve the quality of the data, we filtered SNVs with average read depth less than or equal to 20, or with successful genotype counts less than 7,740 (90%), and subsampled the remaining 233,134 SNVs to 7,740 alleles, which is equivalent to 3,870 diploid individuals (Supporting Information).

## Results

### Comparison with simulated results by FTEC

To validate that the expressions provided in the Methods and Materials section can correctly compute the summary statistics under generalized growth models, we compared the summary statistics calculated by our software EGGS to those simulated by the software FTEC (a coalescent simulator for modeling faster than exponential growth) (Reppell et al. 2012) under the demographic models shown in Figure 2(A). This model is the model of European history inferred in Gazave *et al*. 2014, except that we varied the growth speed parameter *b* (Equation 1), which corresponds to 1 in this model (exponential growth), to also be 0.5 (corresponding to sub-exponential growth), and 1.5 (corresponding to super-exponential growth). The sample size is fixed at 1,000 diploid individuals (2,000 chromosomes). For FTEC simulation, we used mutation rate of 1.2 × 10^−8^ per base pair per generation (e.g., Kong *et al*. 2012) and simulated 200,000 independent loci, each of 1,000 base pairs.

The comparison of the SFS, *S* (across all 200,000 loci), *π* and *α* numerically computed by EGGS to that simulated by FTEC is shown in Figure 2(B)-(E). For each demographic model illustrated in Figure 2(A), the values for all summary statistics from the numerical computation by EGGS are practically identical to those from the simulation results by FTEC. However, our software EGGS exhibits a huge speed improvement over FTEC. For each model considered in Figure 2(A), EGGS takes less than a second to generate the results, while it takes about 5 hours for FTEC to simulate the sequences, due to the large number of independent loci required for accurate estimation (performed in Ubuntu system with Intel Xeon CPU @ 2.67GHz). For instance, when 2,000 independent loci are simulated, which still takes about 3 minutes, the summary statistics deviate considerably from the accurate results (Figure S2 and Table S1). Furthermore, our software works well over a wide range of values of the growth parameter *b*, even when *b* is 0 (corresponding to linear growth or decline) or *b* < 0 (Figure S3), conditions that are not handled by FTEC. We note, however, that as a simulation program FTEC provides the full sequences as output and can have a wider range of applications than facilitated by the SFS and other summary statistics that EGGS calculates.

### Evaluating inference of generalized growth based from the site frequency spectrum

We next set out to test the accuracy (as a function of sample size) of inferring parameters in models with generalized growth from the SFS. A recent study has shown that, in theory, an underlying generalized growth demographic model can be uniquely identified by the ideal, perfect expected SFS of a very small sample size generated from that model (34 haploid sequences for the models shown in Figure 2(A)) (Bhaskar and Song 2014). However, the SFS is estimated in practice from limited amount of data from each individual (even in the case of whole-genome sequencing) and, hence, the estimated SFS will fluctuate around the expected values, which limits its accuracy for inference (Terhorst and Song 2015). We aim to test such inference in practice and determine the power of generalized growth detection and the sample size needed for accurately recovering the growth parameter and other parameters of the demographic model. To be comparable with many practical applications, we considered sequence length that is about equivalent to that obtained from whole exome sequencing (Supporting Information).

We performed inference on the SFS calculated from simulated sequences generated by FTEC. We simulated a demographic model with the same initial epochs as the model illustrated in Figure 2(A). Starting 620 generations ago, the simulated model included a constant population size of 10,000 until 200 generations ago, when the population starts a generalized growth epoch till the present. The generalized growth epoch starts with a population size of 10,000 that grows to an extant effective population size of 1 million individuals, with the growth speed parameter *b* taking either of the following values: 0.4, 0.7, 0.9, 1.0, 1.1, 1.3 and 1.6. We chose these values to represent a range of super-exponential and sub-exponential growth, with emphasis on values around the exponential rate (*b* = 1.0) in order to test the detection power of generalized growth when the growth speed deviates slightly from exponential. We varied the sample size (number of diploid individuals sampled at present) to be 1,000, 2,000, 3,000, 5,000 and 10,000 (Supporting Information). The first 15 entries of the site frequency spectra for these simulated scenarios are shown in Figure S4. From each set of simulations, we then infer four parameters of the recent growth epoch, which can uniquely determine the epoch: 1) the growth speed parameter *b*; 2) the initial population size before growth *N*_i_; 3) the ending population size after growth *N*_f_ ; and 4) the onset time of growth *T*, which is equivalent to the growth duration since the simulated epoch ends at present.

As sample size increases, the accuracy of the point estimates generally improves and the confidence interval narrows (Figure 3). Specifically, when the SFS of only 1,000 diploids is used for inference, the inference performs badly for all parameters, defined by large confidence intervals (Figure 3). However, the confidence interval always includes the true simulated value. A sample size of 2,000 already exhibits acceptable performance except when the growth speed becomes large (*b* = 1.3 and 1.6). Yet larger sample sizes of 5,000 and 10,000 are sufficient for inferring all parameters with very tight confidence intervals. For such sample size, the inference even significantly distinguishes between growth speeds (*b* = 0.9 and *b* = 1.1) that are close to exponential (*b* = 1.0) from that of an exponential, thereby concluding that a sub-exponential (0.9) or super-exponential (1.1) growth has taken place. These observations suggest that a sample size of at least 3,000 diploid individuals might be needed for inferring the parameters associated with the simulated recent generalized growth epoch, which is motivated by previous models of European demographic history. It remains to be explored how accurate are the estimates, and how their accuracy improves with sample size, across a more diverse set of models.

**Figure 3.**
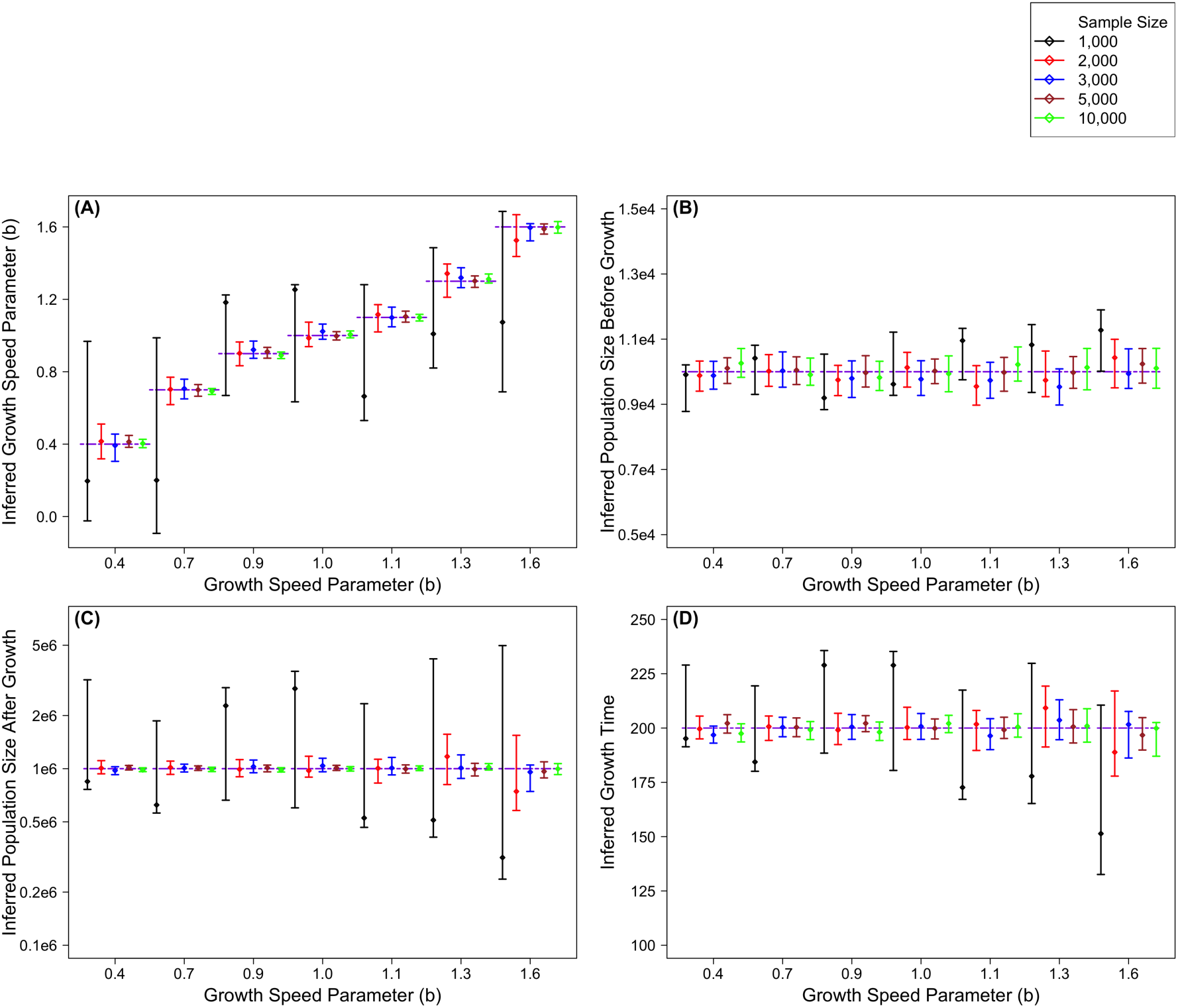
Inference results on simulated data with a recent growth epoch with the following parameters: Growth starts 200 generations before present from an effective population size of 10,000 and ends with an effective population size of 1 million at present. The growth speed parameter *b* takes the following values in different simulations: 0.4, 0.7, 0.9, 1.0, 1.1, 1.3, and 1.6. Inference of these four parameters is based on the SFS estimated from a sample of individuals of one of five sizes (1,000, black; 2,000, red; 3,000, blue; 5,000, dark red; and 10,000, green). The point estimates with 95% confidence interval for these are grouped by the growth speed parameter *b* (*x-axis*). The dashed purple lines show the true values of the simulated model. The results are shown in the following order: (A) the inferred growth speed parameter, (B) the inferred population size before growth, (C) the inferred population size after growth, (D) the inferred growth start time. The *y*-axis in (C) is on log-scale.

### European demographic history inference

We next performed demographic inference on the NHLBI Exome Sequencing Project (ESP) data (Tennessen et al. 2012; Fu et al. 2013). We applied our inference to these data while considering and comparing two models. Both models assume the ancient epochs before 620 generations ago to be the same as those in the model illustrated in Figure 2(A). We infer the parameters only for the most recent epoch, which is one of *generalized* growth in one model, while is limited to *exponential* growth in the other. The parameters for inference are: (1) population size before growth (*N*_i_), (2) population size after growth (*N*_f_), (3) growth onset time (*T*), which is equivalent to the duration of growth, for both models; and (4) only for the generalized growth model, the growth speed parameter (*b*), which is fixed at *b* = 1 for the exponential growth model. The point estimates and 95% confidence intervals are shown in Table 2 and the best-fit demographic models are illustrated in Figure 4.

**Figure 4.**
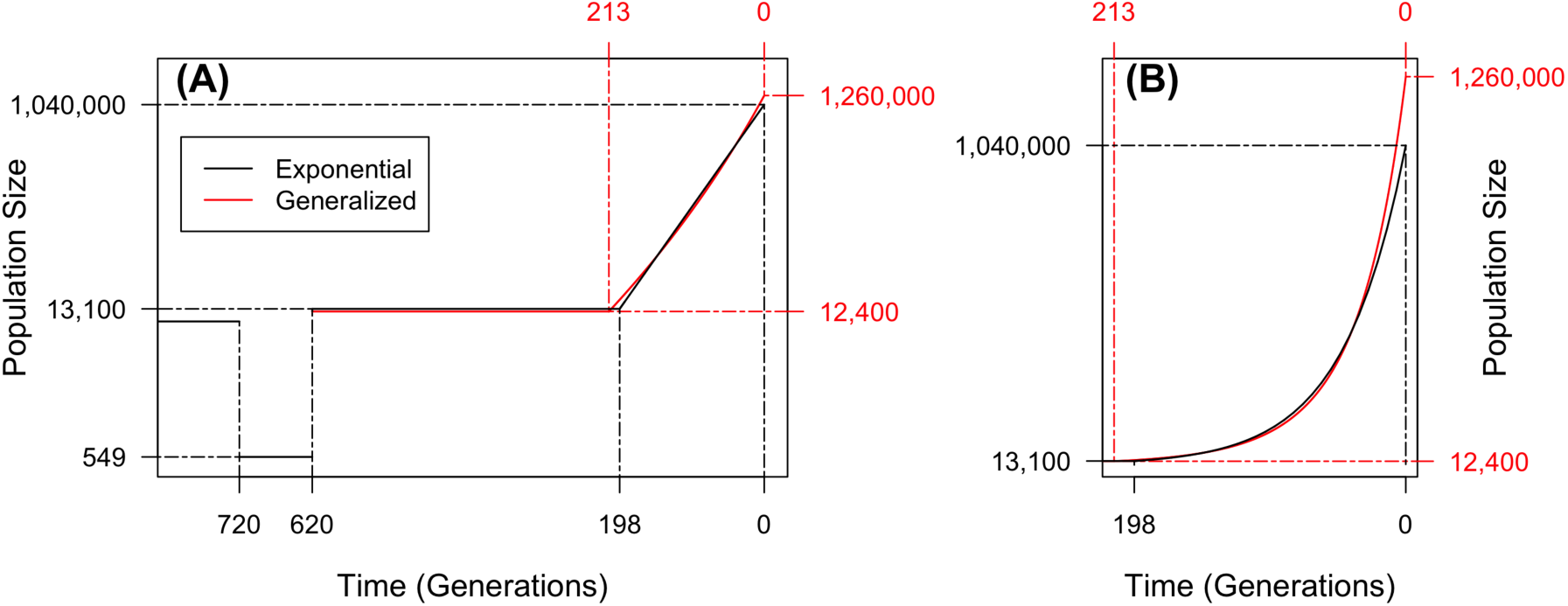
(A) Illustration of the effective population size (*y*-axis, on log scale) over time for the best-fit models inferred based on ESP data. Two models are shown: One restricted to recent growth being exponential (red) and one with a generalized recent growth epoch (black). Before 620 generations ago, the model is not inferred and all parameters are set to be the same as those shown in Figure 2(A). Solid lines show the effective population size over time of each of the inferred models, with dashed line indicating estimating parameter values on the *x*-axis or *y*-axis. Only the most recent 800 generations are shown to emphasize the difference between the two models. (B) A zoom-in to the most recent 200 generations of the inferred models to emphasize the acceleration pattern of the generalized growth model, with *y*-axis on normal scale.

Although our *exponential* growth model assumed a different ancient history before the recent growth epoch than that assumed in Tennessen *et al*. 2012, using ESP data and assuming exponential growth, the inferred growth epoch is generally consistent with that obtained in their study (Figure 4 and Table 1). Our study infers recent growth that started 198 (95% CI: 195-202) generations ago from effective population size of ∼13,100 (12,600-13,600) and continued at a rate of 2.2% (2.15%-2.26%) per generation (Table 2), while Tennessen *et al*. 2012 estimated that recent growth had an initial population size of ∼9,500 individuals, a duration of 204 generations and a growth rate of 2.0% per generation.

**Table 1.** Demographic inference results using ESP data for a model with a recent epoch of *exponential* growth and a model of recent epoch of *generalized* growth. Shown are point estimates and 95% confident intervals (in parenthesis) for the following parameters of the inferred epoch: population size before growth (*N*_i_), population size after growth (*N*_f_), time growth started, in generations (*T*), and the growth speed parameter (*b*), which is fixed at *b* = 1 in the exponential growth case.

| | $N_i (10^4)$ | $N_f (10^6)$ | $T$ | $b$ |
| --- | --- | --- | --- | --- |
| Exponential Growth Model | 1.31<br>(1.26~1.36) | 1.04<br>(1.00~1.07) | 198<br>(195~202) | N/A |
| Generalized Growth Model | 1.24<br>(1.18~1.30) | 1.26<br>(1.16~1.37) | 213<br>(206~220) | 1.12<br>(1.07~1.15) |

The inferred *generalized* growth model fits the data significantly better than that with *exponential* growth (*p*-value = 3.85 × 10^−6^ by *χ*^2^ likelihood-ratio test with 1 degree of freedom). It estimates that growth started 213 (206-220) generations ago from an effective population size of ∼12,400 (11,800-13,000), both values consistent with those estimated in the *exponential* growth model. The extant effective population size following growth is estimated to be 1.26 (1.16-1.37) million. The inferred growth speed parameter *b* = 1.12 (1.07-1.15) is significantly larger than exponential speed of *b* = 1 (*p*-value << 10^−12^ using one-tailed *z*-test), which is the main difference between the two models. *b* = 1.12 implies a growth rate acceleration pattern (Supporting Information) that is super-exponential at 12% faster than exponential through the epoch (Figure 4): the super-exponential growth is relatively slow around the onset time, and it keeps accelerating as time approaches present.

Motivated by these results, we considered a third model with *two* recent exponential growth epochs, which still assumes the ancient epochs before 620 generations ago to be the same as those in the model illustrated in Figure 2(A). Five parameters are inferred (Table S2), with the first phase of growth estimated to have started 219 (95-334) generations ago with a population size of 12,200 (11,700-13,200). This phase of growth lasts until 135 (25-157) generations ago and leads to a population size of 47,100 (30,200-540,900). The population size after the recent phase of growth is 1.12 (1.07-2.09) million. This model provides a significantly better fit than the model with a single *exponential* growth (*p*-value = 5.55 × 10^−6^ by *χ*^2^ likelihood-ratio test with 2 degrees of freedom), but is a worse model than the *generalized* growth model (based on Bayesian Information Criterion, BIC_two-epoch-exponential_ – BIC_generalized_ = 6.1). However, this model exhibits some of the same accelerating pattern as the *generalized* growth model, as ascertained by the growth rate of the most recent exponential epoch being 2.4% (2.3%-5.2%), larger than that of the first exponential epoch, 1.6% (1.3%-2.1%). This acceleration pattern shown in both the generalized model and the model with two exponential epochs is consistent with evidence of growth in European census population size that has greatly accelerated in Modern Era (Keinan and Clark 2012).

## Discussion

In this study, we provide the mathematical derivation and a software that can efficiently compute the expected values of five genetic data summary statistics given a generalized demographic model by evaluating the derived explicit expressions. These summary statistics include the time to the most recent common ancestor (*T*_MRCA_), the total number of segregating sites (*S*), the site frequency spectrum (SFS), the average pairwise difference between chromosomes per site (*π*) and the burden of private mutation (*α*). The fast and accurate generation of these summary statistics is not limited to human demographic inference and will facilitate a variety of population genetic studies in humans and other organisms.

It is also possible that other families of growth models may fit the pattern of human population size history. For instance, previous studies also considered the algebraic-growth model in the form of *N*(*T*) = *T*^γ^ (Eldon et al. 2015). In reality, however, not all demographic models have numerically stable closed-form expressions for the expected time of the first coalescent event (*Ψ*_*j*_). In these cases, fast and accurate numerical integration methods, such as Gauss-Legendre quadrature used in this work, can be applied to evaluate *Ψ*_*j*_. This technique holds the promise of efficiently generating the population genetic summary statistics under arbitrary population size functions.

In future studies, it will be valuable to incorporate gradient-based optimization techniques for the fast inference of demographic models containing generalized growth epochs, e.g. by extending the work of Bhaskar *et al*. 2015. Beyond the direct application for demographic inference and other population genetics analyses, several problems under the framework of Λ-coalescent (Birkner et al. 2013) can also be explored by using the results from this study. If the sample size is large and multi-merger and simultaneous-merger events are permitted, it is shown that under exponential growth models, the expected number of rare variants differs greatly from that derived from Kingman’s coalescent (Bhaskar et al. 2014). This can be extended to the case of generalized growth. For instance, when the growth speed is faster than exponential, the SFS of the Λ-coalescent is expected to differ even more greatly from the of Kingman’s coalescent. Similarly, following the recent work of Eldon *et al*. 2015, a natural question to ask is in what cases can the SFS distinguish between generalized growth models and multi-merger coalescent events.

By applying inference of generalized growth on the SFS generated from the synonymous variants of 4,300 individuals of NHLBI ESP dataset (Tennessen et al. 2012; Fu et al. 2014), we find that generalized growth model shows a better fit to the observed data than the exponential growth model that has been used by all previous demographic modeling studies (*p*-value = 3.85 × 10^−6^). We also find that the European population experiences a recent growth in population size with speed modestly faster than exponential (*b* = 1.12, *p*-value << 10^−12^ for difference from *b* = 1). This result is consistent with previous speculations that human population might have undergone a recent accelerated growth epoch based on the observation of very rare, previously unknown variants in several sequencing studies with large sample sizes (Nelson et al. 2012; Tennessen et al. 2012; Fu et al. 2013). It is also in line with the super-exponential growth in census population size during that time (Keinan and Clark 2012). With the increasing availability of high-quality sequencing data with large sample sizes (Tennessen et al. 2012; Fu et al. 2013; Taylor et al. 2015), more refined and reasonable demographic models can be estimated for different human populations by the application of generalized growth models.

## Acknowledgments

The authors would like to thank Leo Arbiza for helpful comments, Yun S. Song for insightful comments on earlier versions of this manuscript, and Arjun Biddanda for his careful editing of the software manual. This work was supported by National Institutes of Health grants R01GM108805 and R01HG006849, an award from The Ellison Medical Foundation, and an award from The Edward Mallinckrodt, Jr. Foundation. Feng Gao is a Howard Hughes Medical Institute International Student Research fellow.

